# Fast wavefront shaping for two-photon brain imaging with large field of view correction

**DOI:** 10.1101/2021.09.06.459064

**Authors:** Baptiste Blochet, Walther Akemann, Sylvain Gigan, Laurent Bourdieu

## Abstract

*In-vivo* optical imaging with diffraction-limited resolution deep inside scattering biological tissues is obtained by non-linear fluorescence microscopy^1^. Active compensation of tissue-induced aberrations and light scattering through adaptive wavefront correction^2,3^ further extends depth penetration by restoring high resolution at large depth. However, at large depths those corrections are only valid over a very limited field of view within the angular memory effect^4^. To overcome this limitation, we introduce an acousto-optic light modulation technique for fluorescence imaging with simultaneous wavefront correction at pixel scan speed. Biaxial wavefront corrections are first learned by adaptive optimization at multiple locations in the image field. During image acquisition, the learned corrections are then switched on-the-fly according to the position of the excitation focus during the raster scan. The proposed microscope is applied to *in-vivo* transcranial neuron imaging and demonstrates correction of skull-induced aberrations and scattering across large fields of view at 40 kHz data acquisition speed.

## Main text

Adaptive optics (AO) allows the compensation of tissue-induced aberrations in scanning microscopies^2,3^ by taking advantage of the isoplanetism of low order aberration modes^5,6^ at shallow depth over areas of the order of 100 × 100 μm^2^. However, at intermediate depth (≥ one scattering length) a transition occurs between a regime of dominant aberrations to dominant scattering. In this regime, more complex wavefront perturbations occur and, while compensation is possible^7^, corrections are limited to small fields of view^4,8,9^ of the order of the angular memory effect^10^. A fast wavefront shaping method for the compensation of high-order aberrations and scattering for image acquisition over large field of view is thus lacking.

Technically, larger field of view in imaging through scattering media would require AO correction at an update rate equal to the pixel scan rate divided by the size of the memory effect of the medium in pixel units. While fast wavefront shaping based on high-speed spatial light modulators ^11–13^ and hybrid strategies^8,14,15^ was developed to tackle the fast decorrelation time of biological tissues in depth, their update speeds remain too slow to keep up with typical pixel scan rates. Sufficiently fast wavefront update was so far only demonstrated in one dimension and without implementation into a microscope^16^.

Here, we propose a system for adaptive wavefront control at pixel scan speed based on an acousto-optic deflector (AOD) scan engine used in 3D random-address two-photon scanning microscopy^17^. The scan unit consists of a pair of crossed (X,Y) acousto-optic deflector (AOD) units providing free programmable biaxial wavefront correction for every scan point (Fig. 1a, Supplementary Fig. 1). For this, acoustic wave generation in the AODs is synchronized to the arrival of laser pulses from a low repetition rate laser (40 kHz) (Fig. 1a). Pulse-synchronous injection of a frequency modulated acoustic wave generates a spatially varying diffraction grating in each AOD which shapes anew the wavefront of every individual laser pulse. In this way, each AOD acts as a pulse-to-pulse 1D spatial light modulator (SLM) taking control of the local iso-axial slope of the wavefront (Supplementary Fig. 2). Due to the crossed configuration, 2D control is obtained within the limits of corrections linearly separable in X and Y directions^18,19^ (Supplementary Fig. 2 and Supplementary Text). Phase modulation extends to large amplitudes, free of 2π phase jumps, at a resolution of up to 54 orthogonal phase modes per AOD^18^. Finally, combination of AO corrections with tip-tilts and defocus enables 3D image- and random-access scan with synchronous complex wavefront shaping at 40kHz rate (Fig. 1a and Supplementary. Fig. 2).

**Figure 1.**
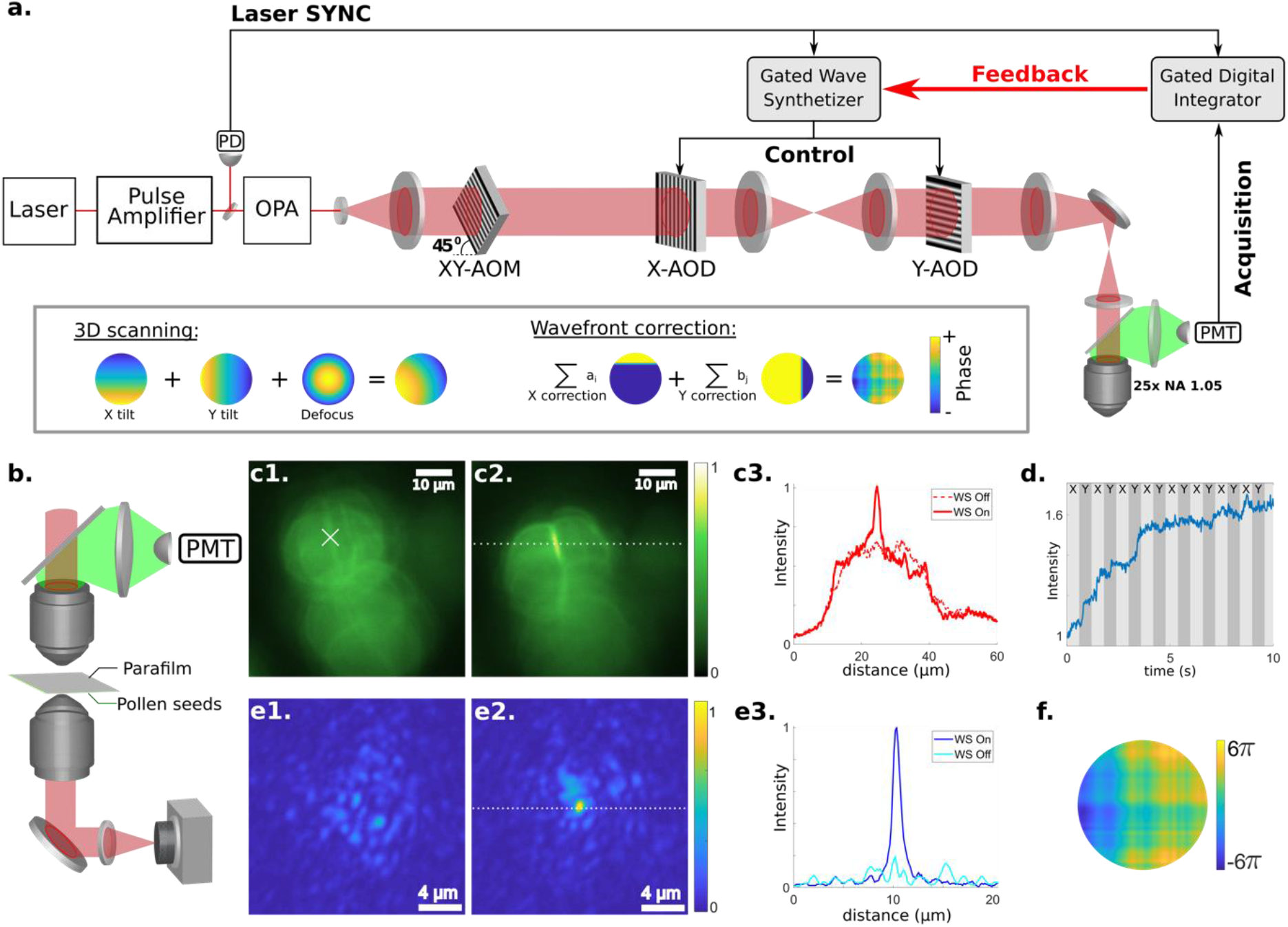
Principle of wavefront optimization in the AOD-based two-photon microscope. **a**. 3Dscope. Photomultiplier (PMT), Photodiode (PD), Optical parametric amplifier (OPA), acousto-optic modulator (AOM), acousto-optic deflector (AOD). Inset: Examples of wavefront engineering for 3D scanning and wavefront correction with two crossed AODs. **b.** Optical setup used to characterize the optimization process. A second microscope objective images the speckle pattern and the focus after optimization for passive monitoring of the optimization. **c.** Two-photon images of pollen seeds positioned behind a scattering medium (two layers of parafilm) before (c1) and after (c2) wavefront correction. Scale bars = 10 μm. c3: Cross section of the two-photon images without (WS off) and with (WS on) wavefront correction. In the isoplanatic patch, wavefront shaping restores resolution and improves the signal intensity. **d.** Evolution of the two-photon signal during the optimization. **e.** Passive monitoring of the point spread function before (e1) and after (e2) wavefront shaping. e3: cross-section of the speckle and of the focus formed without (WS off) and with (WS on) wavefront correction. Scale bars = 4 μm. **f.** Correction phase mask at the end of the optimization.

To find suitable phase corrections, we developed an iterative search algorithm (Supplementary Fig. 3). The correction phase is iteratively constructed from the modes of a chosen 1D basis set (see Supplementary Text) by testing one mode after the other with up to 12 different amplitudes. For this, the mode to be tested is added to the correction mask obtained in the previous iteration and the amplitude maximizing the two-photon feedback signal is retained. The new correction mask is then obtained by adding the tested mode with retained amplitude to the previous correction mask and the iteration proceeds to the next mode until all modes are done. This search is successively performed several times for the modes in X- and Y- direction until convergence, usually after two to five optimization loops. We tested this algorithm first in trans-detection, in which case the feedback signal was the intensity of a single speckle grain collected through a pinhole by a photodiode placed behind a ground glass diffuser (Supplementary Fig. 4). Optimization, at a rate of 5 kHz per mode, generated a well-defined focus on a CCD camera (Supplementary Fig. 4). Notably, the optimization gain (defined as the ratio between the focus intensity after optimization and the mean speckle intensity before optimization) more than doubled when the optimization included X- and Y- instead of only one direction (Supplementary Fig. 4), validating the interest of biaxial AO phase control.

For in-vivo application, the optimization algorithm has to contend with a generally much weaker two-photon fluorescence signal collected in epi-detection as feedback. As a proof of principle experiment, we used fluorescent samples (beads and pollen seeds) placed behind a surrogate scattering medium (parafilm layers) (Fig. 1b and Supplementary Fig. 5). An optimization rate of 33 Hz per mode was achieved by integrating the feedback signal over 100 laser pulses (Supplementary Table 1). Absence of an initial guide star leads to a competition between speckle grains within the same object, until a single speckle grain focus is obtained. Due to this competition, larger size of the fluorescent object required longer optimization time (Supplementary Fig. 5). For small objects such as 2 μm beads, the optimization required less than one second, while for larger objects such as pollen seeds (~20 μm) the optimization lasted up to ten seconds. Images of pollen seeds behind parafilm layers before and after correction (Fig. 1c1-2) showed that high resolution is recovered around the target position within the range of the memory effect (Fig. 1c2-3), with 40 % two-photon signal enhancement (Fig. 1d), in less than 5 s (Fig. 1d). This gain is linked to the formation of a diffracted limited spot with an intensity enhancement of 260 %, as monitored in trans-detection (Fig. 1e1-3). The corresponding correction phase mask in Fig. 1f involves up to 12π phase modulation. As expected, the two-photon gain was larger for small objects (~300 %) and much larger than for correction of system aberration (~10 %, Supplementary Fig. 6).

Wavefront correction *in-vivo* is further impeded by the difficulty for an optimization algorithm to converge in presence of target fluctuations by brain movement^20^ and fluorescence decline by dye photobleaching. To test whether our approach is resilient to these effects, two-photon structural imaging through the thinned skull of anesthetized adult mice was achieved before and after optimization of the wavefront (Fig. 2). Without correction (Fig. 2a1-c1), two-photon fluorescence images of neurons at 35 to 180 μm depth had very low contrast due to skull light scattering^21,22^. Target neurons (indicated by crosses in Fig. 2a2-c2) were selected for wavefront correction. Due to the relatively smaller size of neuron somata (~10 μm), optimizations were faster than for pollen seeds (Fig. 1d) and were achieved in about 3 s (Fig. 2f) with a fluorescence gain of 1.8 ± 0. 6 (N=9, Fig. 2e), visibly increasing resolution close to the target location (Fig. 2a3-c3; notice sharper edges of target neurons) and involving numerous Zernike modes (Fig. 2d). Within the range of angular memory effect, small objects such as dendrites began to appear (Supplementary Fig. 7) after correction. For comparison, we performed the same experiment after craniotomy through a cranial window (Supplementary Fig. 8). As before, correction achieved considerable gain from neuron somata, albeit lower than in transcranial imaging, with the gain linearly increasing with depth up to 600 μm, beyond which the optimization became less efficient, probably due to larger background signal^23^ (Supplementary Fig. 8c).

**Figure 2.**
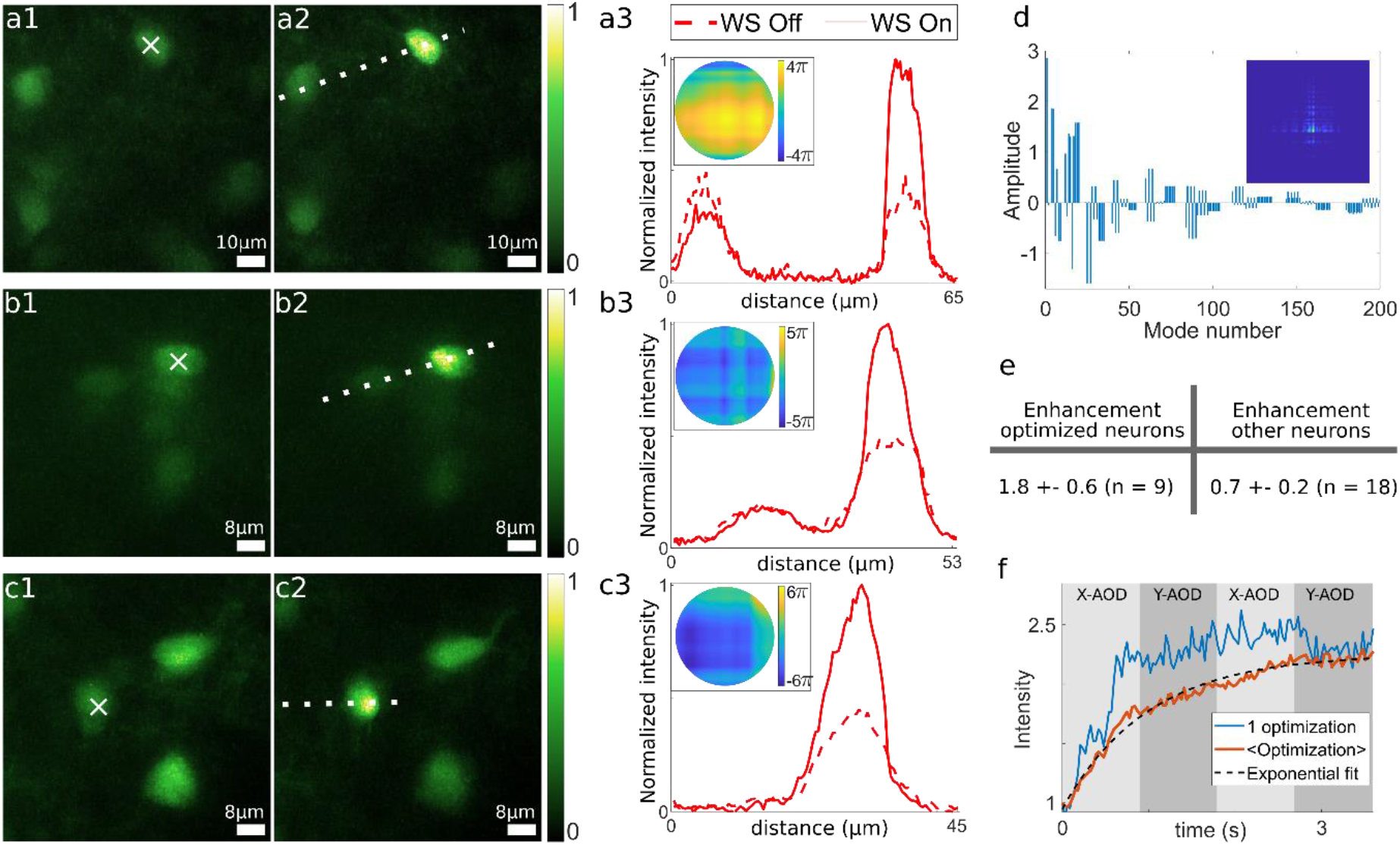
Single-point wavefront shaping correction through thinned mouse skull *in-vivo*. Average skull thickness from 20 μm to 50 μm. **a-c.** Maximum intensity projection of two-photon images obtained before (a1, b1, c1 ) and after(a2, b2, c2) wavefront shaping. Depth below the skull: 90 μm (a), 180 μm (b) and 35 μm (c). Stars indicate the target neurons for the wavefront optimization. Cross-sections of the previous images according to the white lines (a3, b3, and c3). Insets: Corresponding correction phase masks. Images of neurons out of the isoplanatic patches (~15-20 μm) are degraded. Dendrites are visible after wavefront shaping (c3). **d.** Zernike decomposition of the phase mask obtained in b2. Inset: Corrected PSF according to the phase mask obtained in b2. **e.** Summary of the two-photon signal gain obtained inside and outside the isoplanatic patch area (mean ± standard deviation). **f.** Two photon signal during wavefront optimization: single iteration (blue) and average (orange, n=11). Dotted line : exponential fit with a characteristic time of 1.0 ± 0.1s.

However, as shown in Fig. 1c and Fig. 2a2-c2, the area of improved image quality was considerably smaller than the image size. Moreover, outside of this domain, the applied correction downgraded neuron fluorescence intensity by a factor of 0.7 ± 0.2 (N=18) (Fig. 2e and see non-target neurons in Fig. 2a2-c3 and the intensity profile in Fig. 2a3). The correction therefore unfavorably perturbs the local wavefront outside of the isoplanatic range of the angular memory effect in cortical brain tissue of the order of the mean distance between neurons^4^. To overcome this limitation, we devised the search algorithm to retrieve and store adequate corrections for multiple isoplanatic areas of the image field and enabled the scan engine to switch between these corrections on-the-fly during image acquisition according to the focus position during scan progression. The scheme takes advantage of the relatively static nature of the wavefront perturbations introduced by the cranial bone which ensures the corrections to remain valid for prolonged scan times. The benefit of the scheme is apparent in Fig. 3 showing images of two different fields of view (100×100 μm) without correction (Fig. 3a1-b1), with optimization in a single patch and correction applied to the whole image (Fig. 3a2-b2) and with optimization and correction performed in four sub-domains (Fig. 3a3-b3) together with intensity profiles (Fig. 3a4-5 and 3b4-5). As expected, only the last method provided signal enhancement over the entire field of view. For further evidence, we applied the same optimization to other static scattering samples and fluorescent objects yielding similar image enhancement well beyond the size of the angular memory effect of the scattering medium (Supplementary Fig. 9).

**Figure 3.**
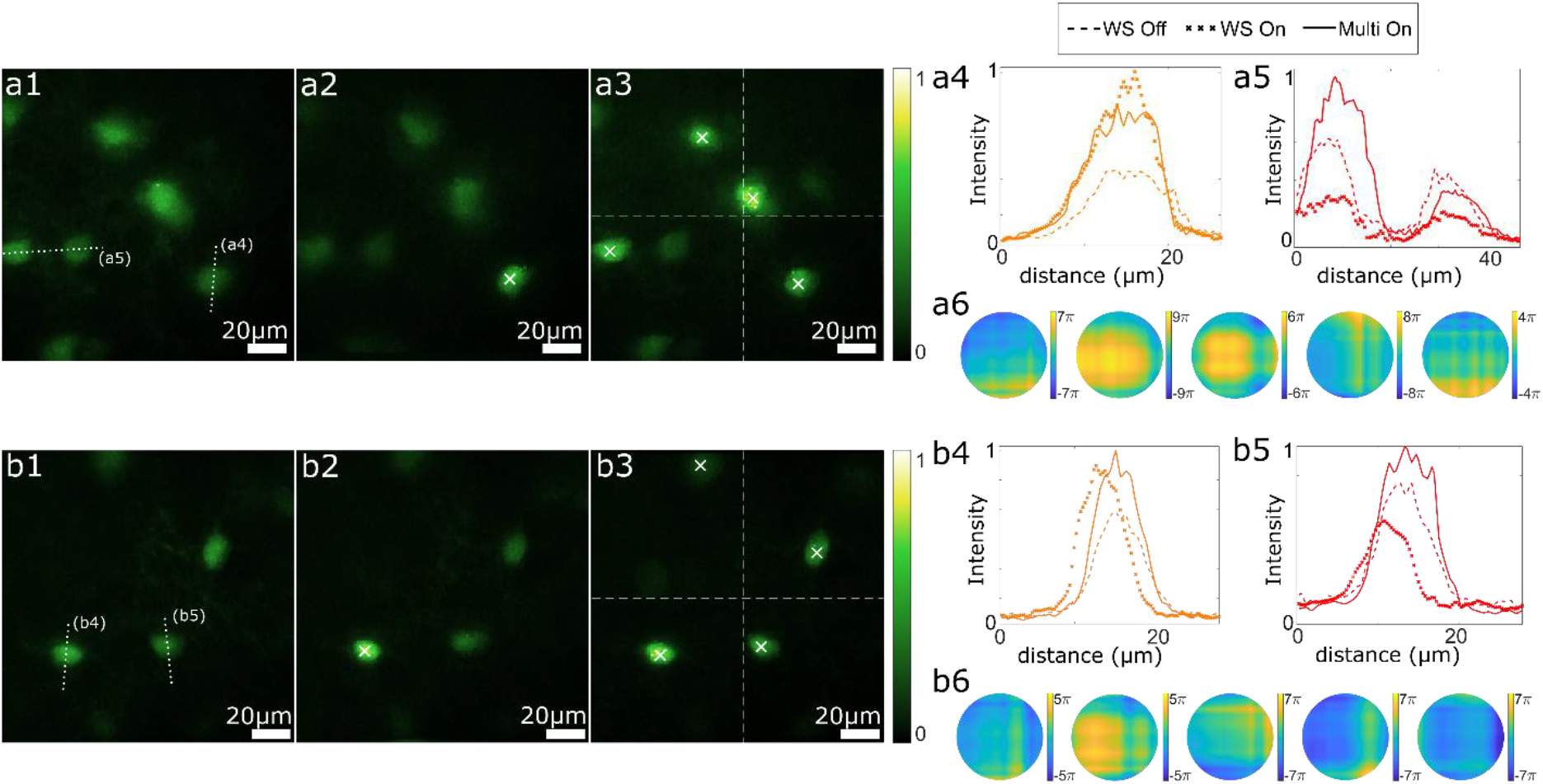
Multi-point wavefront correction through thinned mouse skull *in-vivo*. Average skull thickness about 20-50 μm. **a1-3** and **b1-3**: Maximum intensity projection two-photon images before wavefront correction (a1 and b1), after single correction (a2 and b2) and after multipoint corrections (a3 and b3). Crosses indicate target neurons on which wavefront optimizations were realized. **a4** and **b4**. Cross-sections of target neurons in a3 and b3. Single corrections and multi corrections present similar signal enhancement. **a5** and **b5** Cross-sections on neurons targeted by the multipoint correction in a3 and b3 but not single point correction in a2 and b2. **a6** and **b6**: Wavefront correction phase masks obtained respectively from left to right: single correction (images a2 and b2), multiple corrections (images a3 and b3, respectively top left, top right, bottom left, and bottom right sub-images). Scale bars = 20 μm. Depths below skull: 50 μm (a) and 125 μm (b).

In conclusion, we propose an all-in-one (scanning and correction) wavefront engineering device based on crossed acousto-optics deflectors and implemented in a two-photon microscope. With the ability to shape each individual laser pulse at will, scattering correction can be performed over large area, overcoming the constraint of the small angular memory effect of light scattering in media. We observed higher correction gain in through-skull imaging than in deep imaging through a craniotomy. In the latter case, optimizations were more difficult to achieve because of larger target motions and larger contribution of dynamic scattering. Light scattering by the cranial bone, on the other hand, is stronger and essentially static. Acousto-optic light modulation together with a fast AO optimization algorithm, offers efficient scattering correction over large field of view in non-linear microscopy and holds particular promise for scattering-corrected fast random-access functional recording in cell biology and the neurosciences^18^.

## Methods

### Animal Welfare

Experimental procedures involving mice were conducted in compliance with French and European (2010/63/UE) legislations relative to the protection of animals used for experimental and other scientific purposes. All procedures were approved by the institutional ethics committee “Charles Darwin” registered at the French National Committee of Ethical Reflection on Animal Experimentation under the number 05 (authorization number: APAFIS # 26667).

### Samples

We used four types of fluorescent samples: fluorescent beads (Supplementary Fig. 5 and 9) fluorescent pollen seeds (Fig. 1, Supplementary Fig. 5 and 6), ex vivo fixed brain slices (Supplementary Fig. 9) and in vivo mouse’s brains (Fig. 2 and 3, Supplementary Fig. 7,8 and 9).

#### Fluorescent beads

A solution of 2 μm fluorescent beads (FluoSpheres, Thermofisher) was dropped on a microscope slide and successively covered by a cover slide and a scattering medium: few layers of parafilm (Parafilm) or a mouse’s skull. A parafilm layer has a scattering mean free path l_s_ = 170 μm (at λ = 532nm), a scattering anisotropy g = 0.77 and a thickness of 120 μm ^24^. The parafilm was covered with water for microscope objective immersion. The piece of cranial bone used in Supplementary Fig. 9 had a thickness approximately 100 to 200 μm (with reported values of scattering mean free path^4^ l_s_ below 55μm and of scattering anisotropy^25^ g = 0.9).

#### Pollen seeds

Commercially available pollen seeds slides (Carolina) were placed below different scattering media (parafilm or skull).

#### Ex vivo fixed brain

OneGAD65-EGFP transgenic animal aged 8 weeks was sacrificed by cervical dislocation and the brain extracted and incubated 14h in a solution of 4% paraformaldehyde/PBS and finally rinsed in phosphate buffer solution (PBS). Coronal slices of 200 μm were then cut using a vibratome and stored in PBS. During imaging, the slices were placed in a Petri dish with PBS and immobilized with nylon strings using a homemade harp-like slice anchor.

#### Mouse skull

Two C57BL6J male mice aged 8 weeks were sacrificed by an overdose of Euthasol, after being placed under deep sedation by overdose of isofluorane (5 min at 5% isoflurane in an induction box). Then the skull was extracted and cut to typically 5 mm × 5 mm and stored in PBS.

#### In-vivo imaging through optical windows and with thinned skull

The in-vivo experiments used three female transgenic mice, 6-12 months old, for heterozygous expression of the EGFP gene under the GAD65 promoter. Surgical procedures were performed under deep isoflurane anesthesia supplemented with an analgesic (Buprenorphine, 0.06 mg/Kg). In two animals, a 5 mm craniotomy was performed on the cranial parietal bone of the right hemisphere over primary visual cortex (2.8 posterior and 2.5 mm lateral of Bregma). The craniotomy was then covered with a round cover glass (5 mm, 0.15 ± 0.02 mm thick, Warner Instruments) as a cranial window and sealed with a UV-curable dental cement. To allow proper window and head fixation during experimental sessions, a titan-printed, biometric fixation head post (Ymetry, France) was fixed to the skull with cement. The third animal was prepared for transcranial imaging by skinning the parietal bone over visual cortex with a surgical micro-ball driller until high optical transparency was achieved at estimated 20-50 μm remaining thickness, before mounting the head post. During experimental sessions, the mouse was kept anesthetized under constant flow (0.8 ml/min) of a O_2_/N_2_O mixture (20:80) with 1-1.5 % isoflurane supplied through a custom-designed breathing mask while the mouse was placed on a heating pad (Harvard Instruments) for feedback-controlled maintenance of 37° body temperature. Throughout the sessions, the respiratory rhythm was monitored with a piezoelectric sensor recording respiratory thorax motion.

### 3D-Scope

The experiments employed a self-built two-photon microscope featuring a two-axis, acousto-optic spatial light deflector for ultra-fast amplitude and phase control of the excitation beam^19^ added to a commercial movable objective microscope (MOM, Sutter Instruments)^18^ (Fig. 1a and Supplementary Fig. 1a-b). The output pulses of a 40 kHz regenerative and parametric laser amplifier (Pharos SP6 and Orpheus OPA, Light Conversion) were synchronized to the 40 kHz refresh cycle of two acousto-optic deflector devices (AODs; 15 mm free aperture size; AA Opto-Electronic, France) in crossed X- and Y-orientation such that full aperture Bragg gratings are established in both orientations (X-AOD, Y-AOD) independently for every laser pulse. The AOD devices were designed for large FM modulation bandwidth (36 MHz at 86 MHz carrier frequency) by exploiting large-angle tangential phase match in biaxial TeO_2_ crystals (AA Electronic). In the optical layout, the two AODs were 4f-conjugated to each other and, with a demagnification of 1.5, to the back aperture of the objective (OLYMPUS XLPLan N 25× 1.05 WMP) which was underfilled with 0.66 radial fill factor. Before reaching the X-Y-AOD tandem, the laser beam passes through a third acousto-optic crystal for compensation of temporal and chromatic dispersion of the AOD-diffracted wavefront, in X- and Y-directions, caused by the TeO_2_ bifringence. To achieve dispersion compensation simultaneously for both AODs with a single compensating element, the compensatory crystal is oriented at 45° and driven with a √2-times higher carrier frequency to produce sufficient negative dispersion along both axes to offset the positive dispersion contributed by the X-Y-AODs^26^.

Spatial light modulation is achieved by frequency-modulating the acoustic carrier wave through direct digital synthesizers (DDS) under control of a 80 MHz FPGA-DIO-Board (NI PXIe-7822R, National Instruments) (Fig. 1a and Supplementary Fig. 1c). The gate array (FPGA) hosted a circuit designed to provide real-time frequency command output (23 bit) to the DDS circuits (DDSPA, AA Optoelectronic) in-phase with the AOD laser pulse repetition clock. The command transfer to the DDSs was organized with the help of timing signals that were generated on a digital counter board (NI-PXe 6612, National Instruments). These included two 14 MHz pulse trains (one for each AOD) of 342 pulses, with every pulse gating a single DDS frequency steps during the digital synthesis of the acoustic frequency function along the active coordinate of the AOD aperture. Under these conditions, digital synthesis corresponds to a 323 pixels discretization grid across a 13 mm AOD active aperture. Using this coding space, we defined two basis sets for phase optimization counting 32 and 54 independent modes, respectively (see Supplementary Text) in order to limit jitter^19^ and minimize the optimization time. Every synthesis cycle is triggered by a timing signal which is received from a photodiode measuring the output of the Pharos pulse compressor. A similar gating scheme is reproduced in the detection path which consists of a hybrid Photomultiplier with a 5 mm GaAs cathode (R11322U, Hamamatsu), a low noise 200 MHz voltage amplifier (DHPVA, Femto, Germany) and a 800 MHz/12 bit ADC adapter (NI-5772, National Instruments) together with a 40 MHz FPGA Module (NI-PXIe 7971R, National Instruments). The board runs a circuit that reads the ADC output during a 60 ns time gate in synchrony with the laser clock and organizes the data transfer to the host computer. At the same time, the data are broadcasted in real time to a peer-to-peer (P2P) network to provide real-time feedback to the AOD controller client. The gated acquisition mode was chosen to suppress photomultiplier dark noise during the 25 μs interval between laser pulses. Remaining background light outside the EGFP emission band is suppressed with an emission filter 520/60 (ET520/60, Chroma).

Furthermore, to confirm the principle of acousto-optic wavefront control directly in the excitation path, we set up a transmission microscope composed of a high NA collimator objective (LEICA HC PL APO 40×/1.1 W) on a 5-axis kinematic mount, followed by a 200 mm achromatic tube lens and a camera port. All control and acquisition software were coded in LabVIEW 2015, including the FPGA circuits through the LabVIEW-embedded Vivado 2014.4 compiler (Xilinx).

### Optimization algorithm and experimental protocol

#### 2 photon imaging and wavefront optimization

The two-photon imaging experiments with wavefront optimization were realized in 3 steps:

1. A volumetric two-photon stack (typically 200×200×20 pixels) was acquired with the middle frame identical to the objective focal plane containing the target object (pixel size : 0.5 × 0.5 × 1 μm^3^). For multi-point correction, 4 targets were selected, one for each quarter of the frame. To ensure the gain after wavefront optimization was not due to a change in defocus, the 4 selected targets were initially centered axially on the focal plane.
2. For each target (1 or 4), a continuous wavefront optimization was achieved. At each iteration, a correction mode from a chosen basis set (local-tilt or sinusoidal mode, see Supplementary Text and Supplementary Fig. 2) was selected. N_step_ amplitudes of this mode were successively added to the current correction mask and for each amplitude the two-photon signal integrated over N_p_ pulses was acquired. The tested mode with the amplitude that maximized the two-photon signal was then added to the previous correction mask, which was updated and used for the optimization of the next mode. In total, 64 local tilt modes were used (32 modes for each AOD) for experiments in epi-detection. 108 modes were used for the experiments with the sinusoidal modes in epi detection and the experiment with local tilt in trans detection. When all modes were optimized, another iteration of the algorithm was performed. The full set of modes was usually optimized two or three times. A summary of the optimization parameters can be found in the Supplementary Table 1.
3. A new volumetric two-photon stack was acquired with the correction masks obtained from the optimization algorithm applied. For multipoint correction, the wavefront correction mask is updated on the flight, without delay.

For deep in vivo imaging through a cranial window (Supplementary Fig. 8), an initial phase correction mask was computed at the superficial layer of the cortex. This mask was more likely to correct the system aberration (optical setup and optical window). This mask was then used as the initial phase mask for the experiments at larger depths. For ex-vivo and for non-biological samples, a 2D image rather than a volumetric stack was used to compare image quality before and after correction.

#### Post processing and data analysis

Images were filtered with a Hampel filter to remove outliers and with a bilateral filter to reduce noise. The images from two-photon stacks were registered to correct small plane to plane movements. Finally maximum projection intensity images were computed. The two-photon enhancement was calculated by dividing the integrated two-photon signals over a neuron soma before and after correction.

## Acknowledgements

We wish to acknowledge expert technical help by Astou Tangara for optical engineering, Yvon Cabirou for mechanical engineering, Gérard Parésys, Toufik El Atmani and Lionel Perennes for electronic engineering, the IBENS imaging facility for technical support and the IBENS animal facility for animal care. Funding was provided by H2020 European Research Council (SMARTIES-724473), ANR (ALPINS ANR-15-CE19-0011), Région Île-de-France (DIM Cerveau & Pensée, ALPINS) and the program Investissements d’Avenir launched by the French government and implemented by the ANR, with the references: ANR-10-LABX-54 (Memolife), ANR-11-IDEX-0001-02 (Université PSL) and ANR-10-INSB-04- 01 (France-BioImaging infrastructure).

## Author contributions

LB and SG designed research, BB coded the FPGA circuit and software, WA performed animal surgery, BB and WA performed experiments, BB analyzed data and created figures, BB, WA, SG and LB discussed the results and wrote the manuscript

