## Supplementary materials for "Fast wavefront shaping for two-photon brain imaging with large field of view correction"

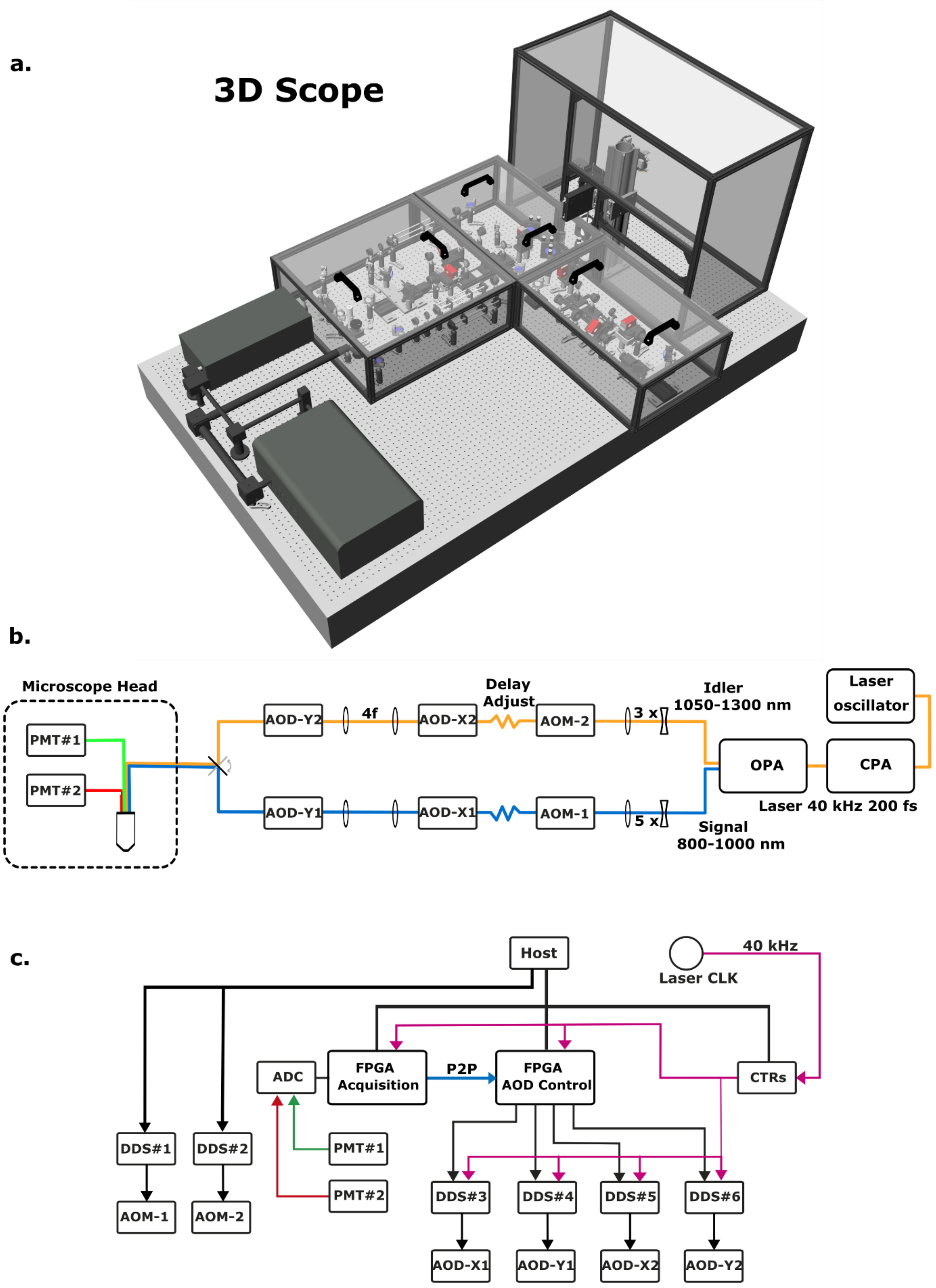


**Supplementary Fig. 1 Microscope setup**

**a.** 3D-rendered view of the microscope (3DScope). **b**. Diagram of the optical beam path featuring two separate acousto-optic deflector units for the signal (blue) and idler (yellow) beams and two-channel detection of fluorescence light in the green (PMT#1) and red (PMT#2) spectral emission range. Only the signal path was used in the present work (see Methods). **c.** Diagram of the electronic layout of the microscope controller. The acquisition and AOD control circuits are implemented on two separate field-programmable gate array (FPGA) boards in a local peer-to-peer (p2p) network for optical phase control under fast feedback (see Methods). Abbreviations: CPA, chirped pulse amplifier, OPA, optical parametric amplifier, AOM, acousto-optical modulator, AOD, acousto-optical deflector with active axis in either X or Y direction, PMT, photomultiplier tube detector, CLK, clock, CTR, Digital counter board, ADC, analog-to-digital converter, DDS, direct digital synthesizer. Connections used to transmit timing signals appear in magenta, p2p connections in blue and analog detector signals in green and red.

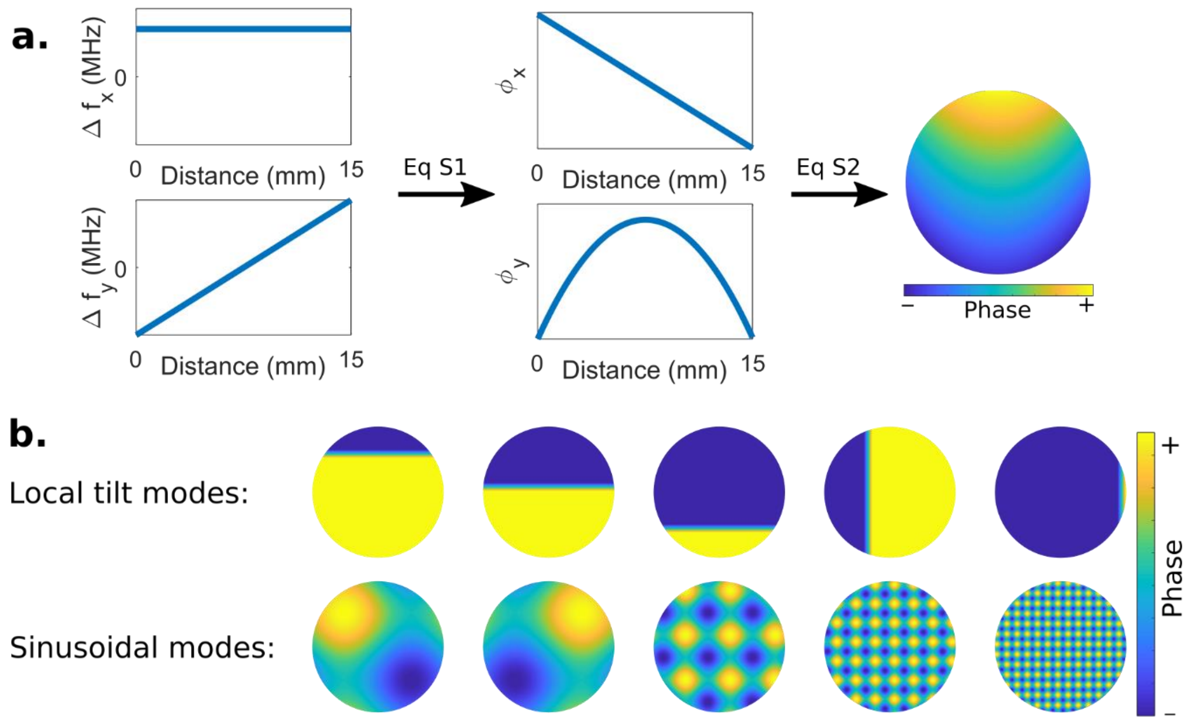


**Supplementary Fig. 2. Wavefront engineering with a pair of AODs.**

**a.** The local frequency of the grating can be tuned at every position *d* within the AOD aperture independently in each AOD. The resulting 1D phase modulation generated in each AOD is given by equation S1 (Supplementary Text-) and the resulting 2D electrical field is given by equation S2 (Supplementary Text). **b.** Examples of set of modes used for wavefront optimization. Local tilt modes (see equations S3 and S4 in the Supplementary Text) were used for the experiments described in the main article and in the supplementary information. The sinusoidal modes (see equation S5 in the Supplementary Text) were used in the experiments described in Supplementary Fig. 5,6 and 8.

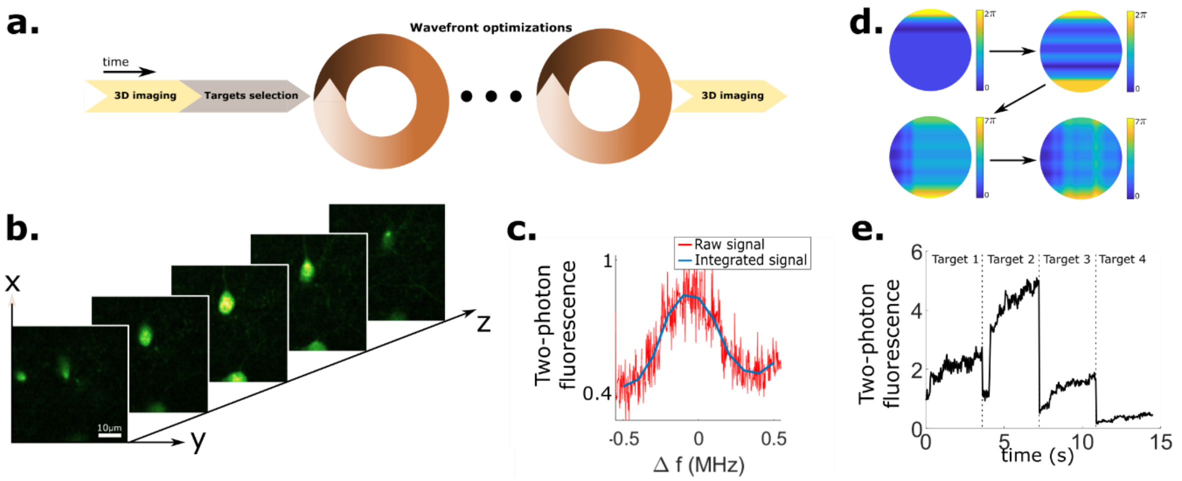


**Supplementary Fig. 3 . Principle of a wavefront optimization experiment.**

**a.** Scheme of the experiment workflow. **b.** A 3D stack is first acquired, and 1 or 4 targets are then selected by the user in the focal plane of the object, which is the middle plane of the stack. There are no constraints on the number of targets. The use of correcting either 1 or 4 targets is shown here by convenience. On each target, an iterative continuous wavefront optimization is realized, and at each iteration the correction phase mask is updated (see Methods). **c.** Example of optimizing the correction mask with regard to a single mode of the basis set: for a given mode, $N_{step}$ amplitudes of this mode are successively added to the correction mask and the two-photon signal is measured for each amplitude (red = raw signal for one laser pulse; blue = signal integrated over $N_p$ laser pulses). A new correction phase mask is finally generated by adding the previous correction mask to the mode just optimized with the amplitude that maximizes the two-photon fluorescence signal. **d**. Examples of successively updated correction masks (order: top-left, top-right, bottom-left, bottom-right) with each iteration optimizing the contribution of a successive mode of the basis set. **e.** Example of the two-photon signal during four consecutive wavefront optimizations on four different targets.

**Supplementary Table 1. Summary of the experimental parameters**

| Experiments | Scattering media | Thickness (µm) | Depth (µm) | $N_{step}$ | $N_p$ | $T_{AODs}$ (s) | $N_{loop}$ | Optimization Time (s) | Power (mW) imaging - optimization |
|---|---|---|---|---|---|---|---|---|---|
| Fig 1 | Parafilm* | 2 layers | | 8 | 100 | 1.2 | 8 | 9.6 | 0.13 – 0.13 |
| Fig 2.a | Skull** | 20-50 | 90 | 12 | 100 | 1.8 | 2 | 3.6 | 0.05 – 0.07 |
| Fig 2.b | Skull** | 20-50 | 180 | 12 | 100 | 1.8 | 2 | 3.6 | 0.23 – 0.35 |
| Fig 2.c | Skull** | 20-50 | 35 | 12 | 100 | 1.8 | 2 | 3.6 | 0.09 – 0.09 |
| Fig 3.a | Skull** | 20-50 | 50 | 12 | 100 | 1.8 | 2 | 3.6 | 0.18 – 0.13 |
| Fig 3.b | Skull** | 20-50 | 125 | 12 | 100 | 1.8 | 2 | 3.6 | 0.13 – 0.10 |

The time to optimize all modes of the 2 AODs once is $T_{AODs} = N_p \times N_{step} \times N_{mode} \times \Delta T$, with ΔT=25µs the inverse of the repetition rate of the laser (40kHz), $N_{step}$ the number of amplitudes successively applied for each mode for search of maximal feedback during optimization, $N_p$ the number of pulses integrated per measurement, and $N_{mode}$ the total number of modes of the used basis set. Typically, $N_p = 100$; $N_{step} = 12$; $N_{mode} = 64$ modes for the 2 AODs. Each AOD was optimized $N_{loop} = 2$ or 3 times before ending the optimization. Note that in all experiments performed in epi-fluorescence involved integration at least $N_p = 100$ laser pulses per feedback measurement, whereas in trans-illumination experiments (Supplementary Fig. 4) $N_p = 1$ because of the larger amplitude of the feedback in this configuration resulting in much faster optimization time.

*: ls(parafilm) = 170 µm; g(parafilm) = 0.77 at λ = 532 nm [1].

**: ls(skull) < 55 µm ; g(skull) = 0.9 at 900 nm [2,3].

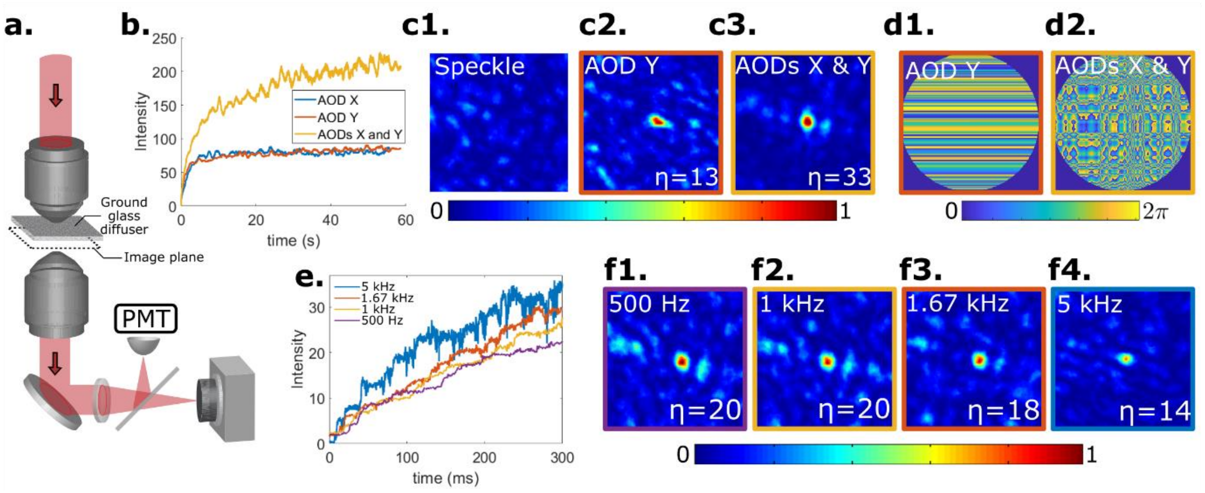


**Supplementary Fig. 4. Optimization in transmission. a.** Scheme of the optical setup. A scattering media (ground glass diffuser) is positioned between the image plane and the first microscope objective. A second microscope objective images the speckle pattern simultaneously on a CCD camera (Basler cA2040-55um) and on a PMT. The PMT collects one speckle grain as a feedback signal for the wavefront optimization. **b.** Comparison between wavefront optimization with one AOD (blue : AOD X and red curve AOD Y) versus two AODs. Each AOD allows controlling 54 modes, with a single-mode optimization rate of 500 Hz. Combined optimization using two AODs offers more degrees of freedom resulting in two times larger final enhancement. **c1**, **c2** and **c3.** Images respectively of the speckle before optimization, of the focus after optimization with 1 AOD and of the focus after optimization with 2 AODs. The enhancement (gain) η is defined as the ratio between the focus intensity after optimization and the mean speckle intensity before optimization. **d1** and **d2.** Correction phase masks obtained respectively after one AOD optimization and two AODs optimization. **e.** Comparison of enhancement for different single mode optimization rate. The fastest single mode optimization rate achieved is 5 kHz (blue curve), which is close to the best rate achieved in the literature for a continuous optimization[4]. **f1**, **f2**, **f3** and **f4**. Images of the focus obtained after 1000 iterations for different single mode optimization rate: 500 Hz (f1), 1 kHz (f2), 1.67 kHz (f3), and 5 kHz (f4). Using fast optimization rate resulted in faster convergence but suffered from lower SNR, which reduced the final enhancement.

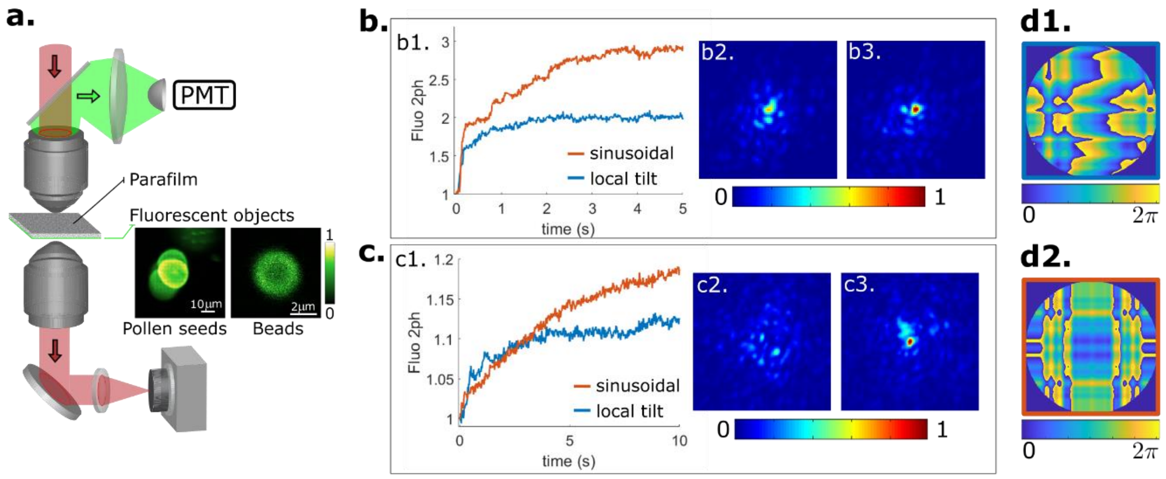


**Supplementary Fig. 5. Optimization in epi-detection**: **Impact of the fluorescent objects size and of the mode basis**.

**a.** Fluorescent objects (pollen seeds or 2 µm beads) are placed behind a scattering medium (2 layers of parafilm). The total two-photon fluorescence collected in epi-detection is used as a feedback for the wavefront optimization. In transmission, a camera (Basler cA2040-55um) monitors passively the focus before and after correction. Insets: two-photon images of the fluorescent objects without scattering medium. **b.** Wavefront optimization with a small guide star (2 µm bead) for two sets of modes: local tilt in blue and sinusoidal in orange (see Supplementary Fig. 2 for more details). **c.** Wavefront optimization with a large guide star (~20 µm pollen seeds) for two sets of modes. **b1.** and **c1.** Average evolution of the two-photon signal during optimization (average over 20 realizations). The sinusoidal basis being smoother allows to control more modes (54 versus 32 in case of the local tilt bas) at comparable error budget in the AOD generated phase mask, which leads to a higher enhancement. . However, the local tilt basis converges faster at short times. Due to competition between speckle grains within the fluorescent object, optimization is longer for extended objects. Enhancement is also smaller due to the signal emitted by the neighboring speckle grains. **b2. c2. b3** and **c3.** Images of the focus measured in transmission before optimization (b2. and c2.) and after optimization (b3 and c3). **d1** and **d2** Examples of correction phase masks obtained respectively with the local tilt basis and the sinusoidal basis.

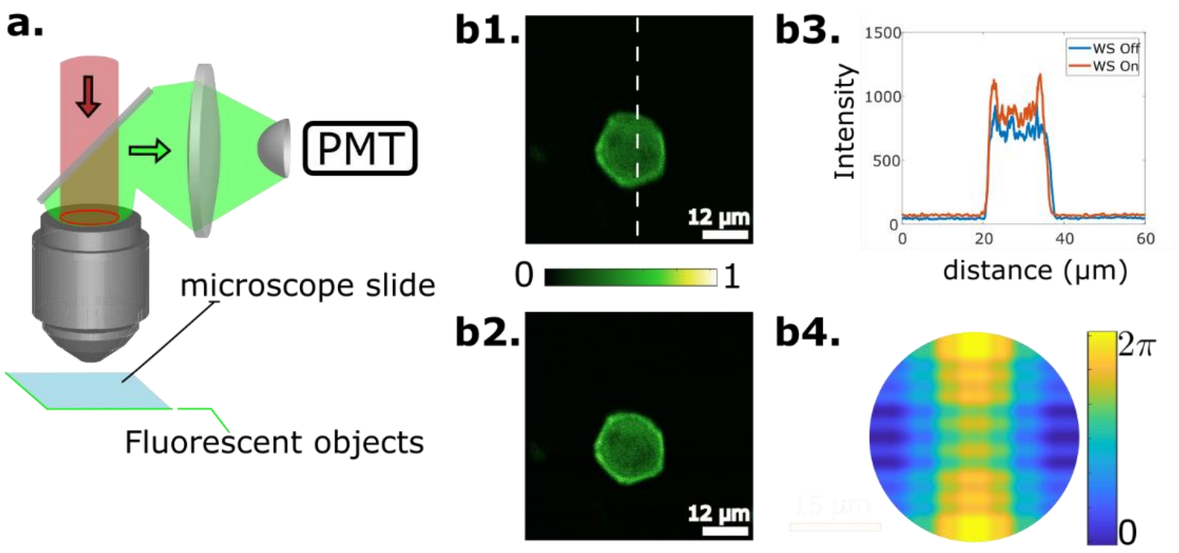


**Supplementary Fig. 6. Optimization in epi-detection: correction of system aberrations.**

**a**. Scheme of the optical setup. **b.** System correction with the AODs using pollen seed as test object. **b1.** Two-photon image before correction. **b2** Two-photon image after correction. **b3.** Cross sections along the dotted line shown in figure b1. **b4.** Correction phase mask.

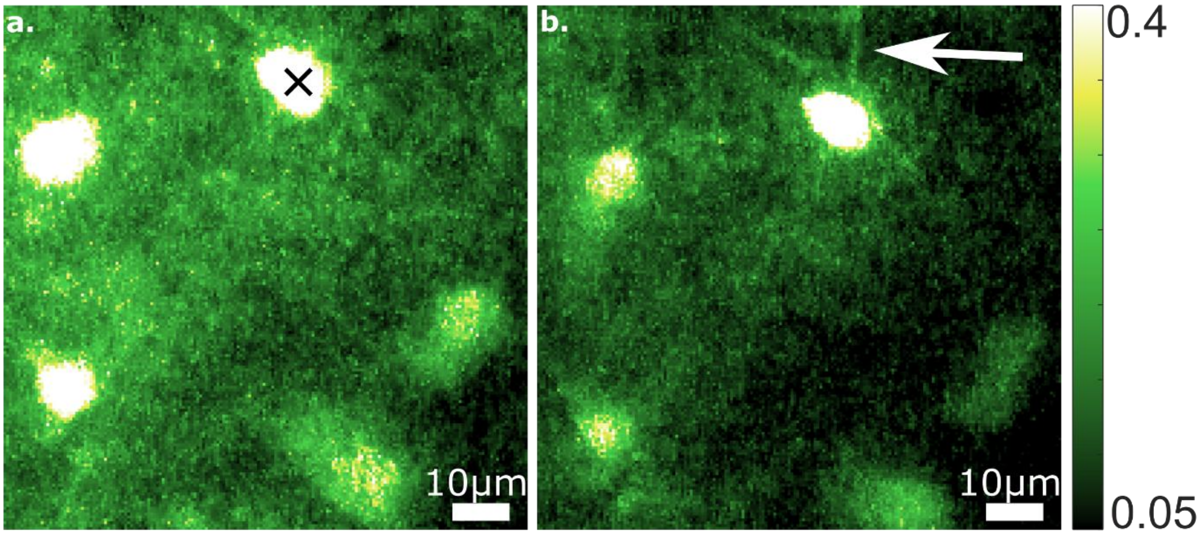


**Supplementary Fig. 7. Image improvement on dendrites through a thinned skull.**

Average skull thickness about 20-50 µm. Saturated maximum intensity projection around proximal dendrites before (**a**) and after (**b**) wavefront correction at 90 µm depth below the thinned skull. The black cross in figure (a) indicates where the wavefront optimization was achieved. Inside the isoplanatic area, dendrites start to become visible after wavefront correction.

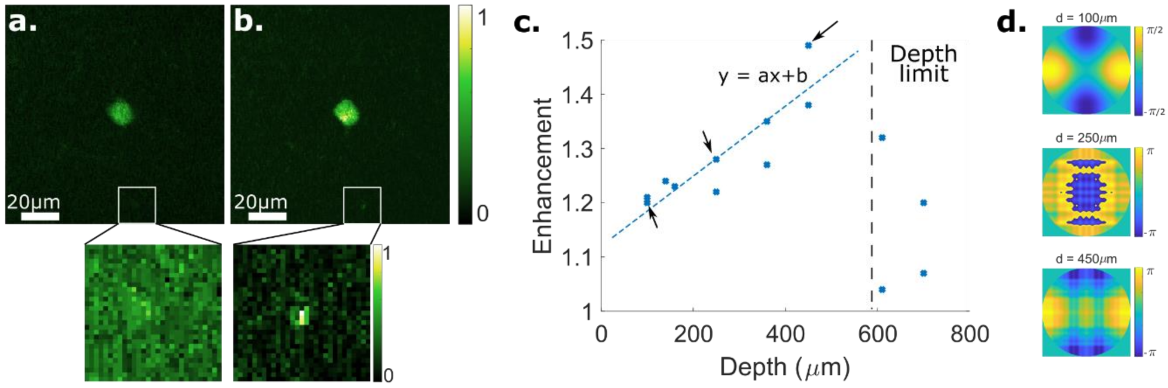


**Supplementary Fig. 8. Wavefront correction in depth through an optical window.**

**a.** and **b.** Maximum intensity projection before (a) and after correction (b) of a neuron imaged at z = 450 µm in the mouse brain. Insets: zooms on a small dendrite branch. **c.** Two-photon enhancement measured as a function of the depth. Dotted line: linear fit to the data. At low and shallow depth, a linear relationship exists between depth and enhancement: y = ax + b with a = 0.6 ± 0.2 $10^{-3}$ $um^{-1}$ and b = 1.1 ± 0.1. At larger depth (> 600 µm), two-photon microscopy starts to fail due to increased fluorescent background. **d.** Correction phases masks obtained for experiments indicated by arrows in c.

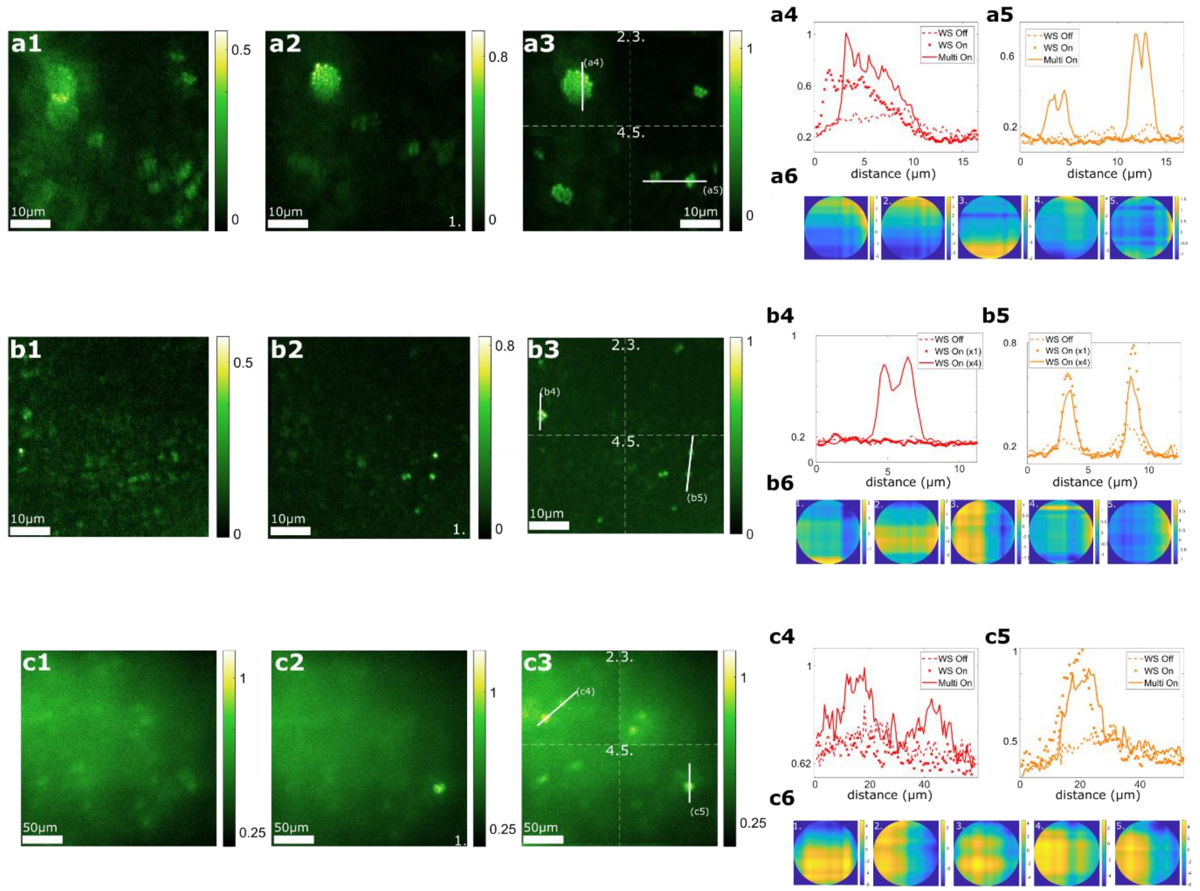


**Supplementary Fig. 9**: **Additional examples of multipoint corrections.**

**a.** Fluorescent beads behind three parafilm layers (different color bars) ; **b**. Fluorescent beads behind a skull (different color bars) ; **c.** a fixed brain slice behind a skull (same color bars). From left to right: Two-photon image without correction (a1,b1,c1) ; two-photon image with correction on one target (a2,b2,c2); two-photon image with corrections on four targets (a3,b3,c3); corresponding cross sections (a4-5, b4-5, c4-5). The corresponding correction phase masks are shown for a single point correction (left) and for multiple points corrections (from left to right field of view labeled 2 to 5).

**Supplementary text.**

The phase pattern $\varphi_{x\ or\ y}$ generated by an $AOD_{x\ or\ y}$ as a function of the local frequency of the acoustic pattern $f_{x\ or\ y}(u)$ at the radial coordinate d of the $AOD_{x\ or\ y}$ aperture is:

$$\varphi_{x\ or\ y}(d) = \frac{2\pi}{v}\int_{-\frac{L}{2}}^{d} \Delta f_{x\ or\ y}(u)du \qquad \text{(S1)}$$

with $\Delta f_{x\ or\ y}(u) = f_{x\ or\ y}(u) - f_c$, the frequency modulation signal, $f_c$, the central frequency, v, the acoustic wave velocity and L, the diameter of the AOD active aperture.

*Local tilt basis set used for phase optimization.* The 2D optical field generated by two cross AODs in a plane optically conjugated to the two AODs is:

$$E(x,y) = e^{i(\varphi_x(x)+\varphi_y(y))} \qquad \text{(S2)}$$

The $i^{th}$ local tilt mode on x or y used for wavefront optimization is defined by:

$f_{x\ or\ y}^{i}(x\ or\ y)$ = 1 for (x or y) ∈ [i-1,i]*2*L/$N_{modes}$.

= 0 elsewhere (S3)

The associated phase patterns are obtained by integrating the function S3 according to equation S1:

$\varphi_{x\ or\ y}^{i}(x\ or\ y)$ = 0 for (x or y) < (i-1)*2*L/$N_{mode}$.

= x*$N_{mode}$/(2*L) for (x or y) ∈ [i-1,i]*2*L/$N_{mode}$

= 1 elsewhere (S4)

with L the diameter of the AOD active aperture.

*Sinusoidal basis set.* The sinusoidal modes used for wavefront optimization are defined by:

$f_{\pm}^{i}(x) = \pm cos(\omega_i.x/L)$ and $f_{\pm}^{i}(y) = \pm cos(\omega_i.y/L)$ (S5)

with $\omega_i$ ranging from 1 to 15. The associated phase patterns are obtained by integrating the function S5 using S1 as before.
